# A high sugar diet, but not obesity, reduces female fertility in *Drosophila melanogaster*

**DOI:** 10.1101/2023.03.02.530894

**Authors:** Rodrigo Dutra Nunes, Daniela Drummond-Barbosa

## Abstract

Many studies from *Drosophila* to humans show a strong link between obesity and reduced fertility. However, obesity is often induced by changes in diet or eating behavior, such that it remains unclear whether reduced fertility is a consequence of obesity itself, diet, or both. Here, we report that a high sugar diet reduces *Drosophila* female fertility by increasing death of early germline cysts (prior to follicle formation) and degeneration of vitellogenic follicles; that obesity in and of itself does not impair fertility; and that high glucose levels closely correlate with reduced fertility on a high sugar diet. Females on a high sugar diet rapidly develop obesity (and display high glycogen, glucose, and trehalose levels, and insulin resistance) and decreased fertility. In stark contrast to high-sugar-obese females, females in which similar levels of obesity are induced by adipocyte-specific knockdown of anti-obesity genes *brummer* or *adipose* have normal fertility and sugar metabolic indicators. Remarkably, females on a high sugar diet supplemented with a separate source of water also have normal fertility and glucose levels, despite persistent obesity, high glycogen and trehalose levels, and insulin resistance markers. These results strengthen our conclusion that obesity itself does not impair fertility, show an inverse correlation between high glucose levels and fertility, and demonstrate that insulin signaling levels remain sufficiently high to maintain insulin-dependent processes during oogenesis irrespective of insulin resistance markers.

## INTRODUCTION

The global obesity epidemic is a major public health concern (www.who.int), and epidemiological and animal model studies show a link between obesity and many negative health outcomes, including loss of fertility [1,2]. In humans, the sharp increase in obesity incidence (largely owing to unhealthy diets and lack of physical activity) correlates with a corresponding decrease in fertility [3]. In mammalian models showing that obesity lowers fertility through various proposed mechanisms (e.g., altered gonadotropin responses, reduced number of eggs and rates of fertilization), obese mice are generated through obesogenic diets or genetic mutations that alter hormonal regulation and eating behavior [4]. *Drosophila* females fed a diet with high sugar levels are also obese and have reduced fertility [5,6]. These studies in mammals and *Drosophila* have thus not distinguished whether effects on fertility are a consequence of the diet itself, the obesity (caused by the diet), or a combination thereof. This major shortcoming has limited our ability to fully investigate the root cause of the fertility decline often attributed to obesity.

The genetic model organism *Drosophila melanogaster* represents a powerful system for investigating fundamental aspects of metabolism and physiology [7] implicated in obesity. *Drosophila* adipocytes reside alongside hepatocyte-like oenocytes in an organ called the fat body [7,8], and lipid metabolism is controlled by evolutionarily conserved metabolic pathways [7]. For example, mutation of the *Drosophila* anti-obesogenic genes *brummer* (encodes a triglyceride lipase) or *adipose* (encodes a WD40/TPR-domain protein) leads to obesity – as does mutation of their mouse/human homologs *Atgl*/*ATGL* and *Wdtc1*/*WDTC1*, respectively [9–13]. Similar to effects observed in mice and humans [9,11], feeding *Drosophila* a diet high in sugar leads to obesity, insulin resistance, hyperglycemia, and cardiovascular abnormalities [6,14,15], further underscoring the metabolic and physiological similarities between *Drosophila* and mammals.

The physiological regulation of *Drosophila* oogenesis has been extensively studied [16]. *Drosophila* females have a pair of ovaries, each subdivided into ∼15 ovarioles (Fig 1A). Each ovariole has a germarium followed by progressively more developed follicles (Fig 1B). Two-to-three germline stem cells reside within a specialized niche (composed primarily of cap cells) in the anterior portion of each germarium (Fig 1C). Each germline stem cell division generates a germline stem cell and a posteriorly displaced cystoblast that undergoes four rounds of incomplete mitoses to generate a 16-cell cyst. Follicle cells surround the 16-cell germline cyst to form a new follicle that leaves the germarium. Follicles develop through 14 recognizable stages of oogenesis to form a mature egg, with vitellogenesis (i.e. yolk uptake) beginning at stage 8. Previous studies on the effects of dietary yeast and diet-dependent signaling pathways identified major points of physiological regulation of oogenesis [16]. For example, insulin signaling in the ovary and adipocytes is required for normal germline stem cell maintenance and proliferation, early germline cyst survival, follicle growth, and progression through vitellogenesis [17–19]. More recently, two groups reported that *Drosophila* females maintained on high sucrose food for seven days are obese, have insulin resistance, and lay fewer eggs than those on low sucrose [5,6]. One study reported that ovaries are smaller on high sucrose [5]. However, two important points remain unclear: which specific stages of oogenesis are affected in obese *Drosophila* females on a diet high in sucrose and how diet versus obesity contribute to their reduced fertility.

**Fig 1.**
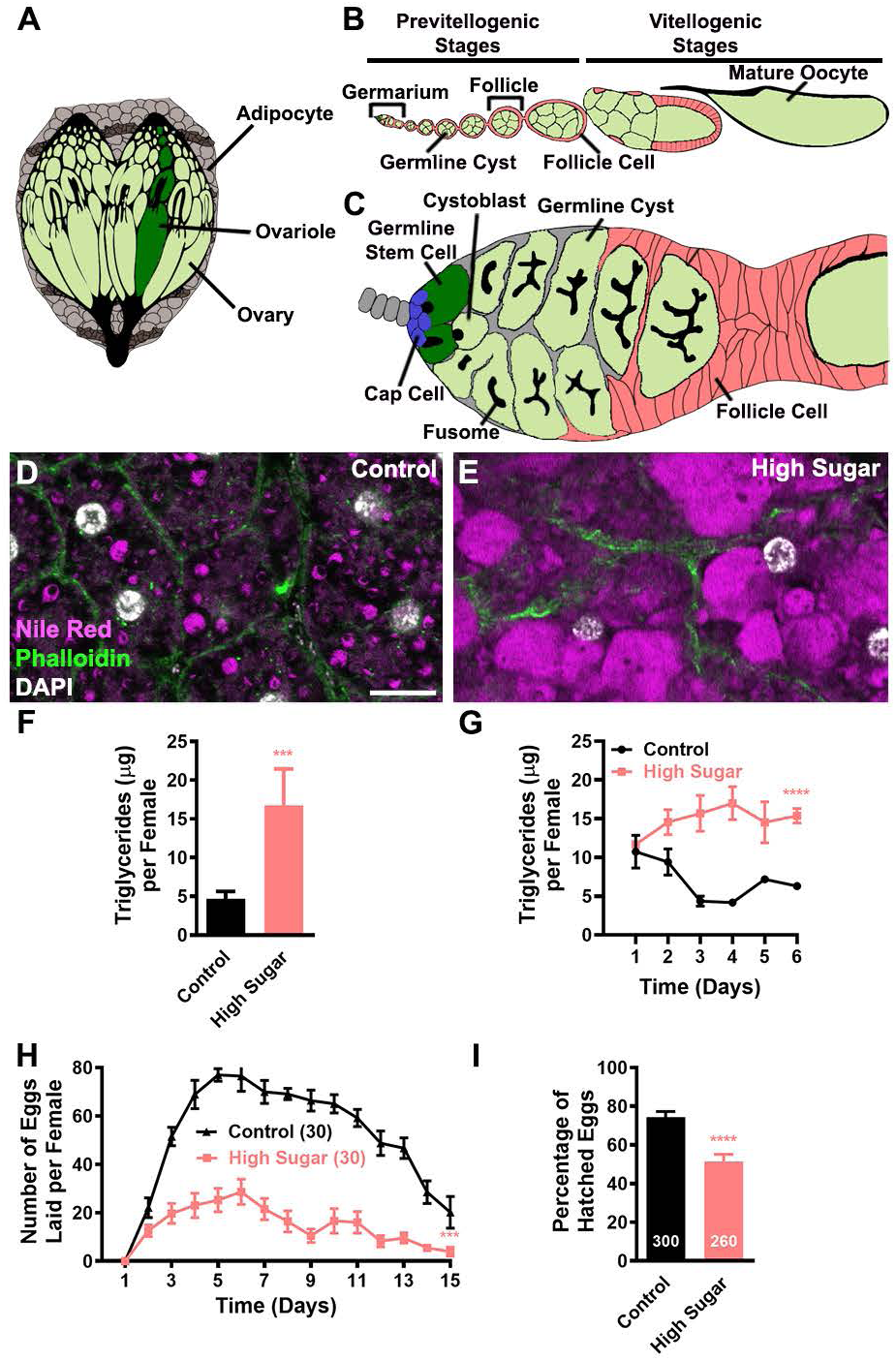
Females on a high sugar diet rapidly develop obesity and produce fewer eggs with reduced hatching rates. (A) Diagram showing a pair of *Drosophila* ovaries (green) near the fat body, which is composed of adipocytes (light grey) and oenocytes (dark grey). Each ovary has ∼15 ovarioles, one of which is highlighted in dark green. (B) Ovariole diagram showing the anterior germarium followed by developing follicles (or egg chambers) that give rise to mature oocytes. Each follicle contains a germline cyst (green; one posterior oocyte plus 15 nurse cells) surrounded by follicle cells (pink). Vitellogenic stages are marked by the presence of yolk in the oocyte. (C) Germarium diagram showing germline stem cells (dark green) in close association with somatic cap cells (blue). Each germline stem cell division renews the germline stem cell and produces a cystoblast that undergoes four rounds of mitosis to form a 16-cell cyst. Follicle cells envelop the 16-cell cyst to form a new follicle that leaves the germarium. (D, E) Adipocytes from seven-day-old females on control (D) or high sugar (E) diets. Nile red (magenta) labels lipid droplets; Phalloidin (green) outlines cell membranes; DAPI (white) labels nuclei. Images represent projections of three 1-μm confocal sections. Scale bar, 10 μm. (F) Bar graph showing average triglyceride content per female (after subtraction of ovarian triglyceride content) of seven-day-old females on control or high sugar diets. Data shown as mean±s.e.m. from three experiments (10 females per condition per experiment). \*\*\**P*<0.001, Student’s *t*-test. (G) Line graph showing average triglyceride content of females on control or high sugar diets over six days. Data shown as mean±s.e.m. from three biological replicates. \*\*\*\**P*<0.0001, two-way ANOVA with interaction. (H) Average number of eggs laid daily per female on control or high sugar diets. The number of females (paired with males) analyzed are shown in parentheses. Data shown as mean±s.e.m. from three experiments. \*\*\**P*<0.001, *F*-test of third order polynomial fitted curves. (I) Average percentage of eggs laid by females on different diets that hatch into larvae. The number of eggs analyzed are shown inside bars. Data shown as mean±s.e.m. from three experiments. \*\*\*\**P*<0.0001, Student’s *t*-test.

Here, we show that adult *Drosophila* females on a high sugar diet become obese and produce fewer eggs as a result of increased death of early germline cysts and degeneration of vitellogenic follicles, and that laid eggs have lower hatching rates. By contrast, no negative effects on oogenesis result from obesity caused by adult adipocyte-specific knockdown of conserved anti-obesogenic genes *brummer* or *adipose*, demonstrating that obesity is not sufficient to decrease female fertility. High-sugar-obese females, but not *brummer*- or *adipose*-knockdown females have increased levels of glycogen, trehalose and glucose, as well as high insulin resistance in fat bodies and ovaries. Intriguingly, when an extra source of dietary water is provided to females on a high sugar diet, they remain obese and maintain high levels of trehalose and insulin resistance markers, yet their egg production rates are largely restored and strongly correlate with a dramatic reversal of elevated glucose levels. Altogether, our findings show that obesity, high glycogen, and insulin resistance are not responsible for the reduced fertility of females on a high sugar diet, indicating that insulin signaling remains above the threshold required for insulin-dependent oogenesis processes. We further show that dietary water-dependent factor(s) (e.g. glucose and/or unknown molecules) are instead responsible for impairing female fertility on a high sugar diet, providing a foundation for future research on the molecular mechanisms linking high levels of dietary sugars to disruption of specific processes during oogenesis.

## RESULTS

### Females maintained on a high sugar diet are obese and produce fewer eggs with reduced hatching rates

Previous studies showed that *Drosophila* females on a high sugar diet are obese and produce fewer eggs [5,20]; we therefore set out to pinpoint what specific stages of oogenesis are sensitive to high dietary sugar. We first confirmed that females fed a high sugar diet become obese and lay fewer eggs under our experimental conditions (Fig 1D-G). Based on Nile red staining of lipid droplets in whole mount adipocytes (Fig 1D-E) and biochemical measurements of fat body triglycerides (Fig 1F), we observed that females maintained on a high sugar diet for seven days had markedly larger lipid droplets and three-fold higher triglyceride levels compared to females on a control diet (Fig 1D-F). To determine how quickly females develop obesity on a high sugar diet, we measured their triglyceride content daily from one to six days of age on control versus high sugar diets. Remarkably, while control females gradually lost triglycerides during their first three days of adulthood (presumably owing to the elimination of larval fat body cells present at eclosion [21]), females on a high sugar diet displayed increased triglyceride content starting within two days of eclosion (Fig 1G). We next measured the number of eggs produced by these high-sugar-obese females daily over a 15-day period. From the earliest time point through the end of the time course, high-sugar-obese females consistently laid fewer eggs compared to females on the control diet (Fig 1H). Moreover, the eggs laid by high-sugar-obese females hatched at a ∼30% lower rate relative to control eggs (Fig 1I), in apparent contrast to a previous study showing no effect of a high sugar diet on hatching rates [5] (see Methods for details). Altogether, these experiments confirm that females on a high sugar diet are obese and less fertile, and further show that obesity and loss of fertility develop concomitantly starting within a couple of days on high sugar.

### Germline stem cell maintenance, germline stem cell proliferation, and follicle growth are not affected by a high sugar diet

We then investigated if and how a high sugar diet affects specific stages and processes of oogenesis known to be regulated by organismal physiology [16], starting with germline stem cells. Germline stem cell numbers in females on control and high sugar diets remained similar at zero, five, 10 and 15 days (Fig 2A and 2B). Accordingly, cap cell numbers were indistinguishable regardless of diet (Fig 2A and 2C). There were no significant differences in the frequencies of germline stem cells labeled by EdU (an S phase marker) (Fig 2D and 2E) or phosphorylated histone H3 (an M phase marker) (Fig 2F and 2G). These results indicate that a high sugar diet has no effect on either maintenance or proliferation of germline stem cells.

**Fig 2.**
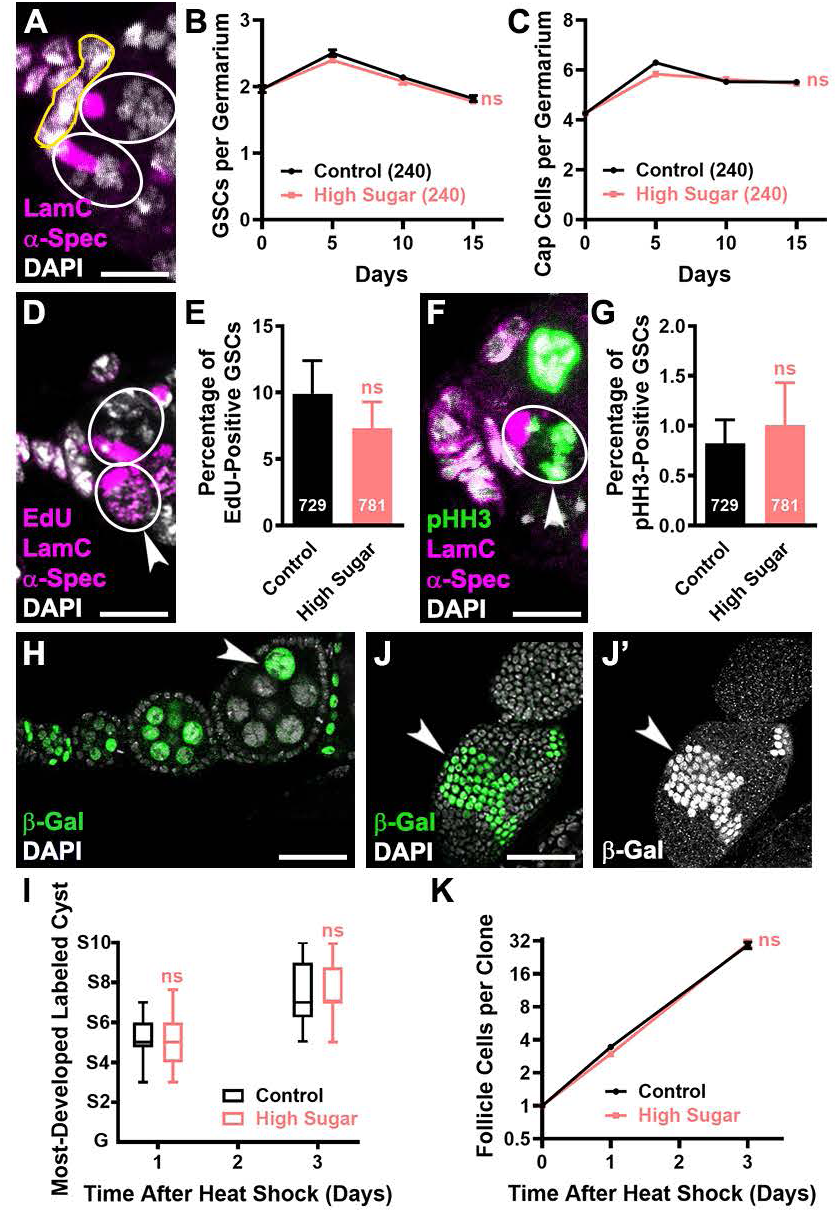
A high sugar diet does not affect germline stem cell maintenance or proliferation or follicle growth. (A) Anterior portion of germarium from 10-day-old control female showing germline stem cells (white outlines) and cap cells (yellow outline). α-Spectrin (magenta) labels fusomes; Lamin C (magenta) labels cap cell nuclear envelopes; DAPI (white) labels nuclei. Image represents a projection of three 1-μm confocal sections. Scale bar, 10 μm. (B, C) Line graphs showing average number of germline stem cells (GSCs; B) or cap cells (C) per germarium in zero, five-, 10-, and 15-day old females on control or high sugar diets. Numbers of germaria analyzed are shown in parentheses. Data shown as mean±s.e.m. from three independent experiments. ns, no significant difference, two-way ANOVA with interaction. (D-G) Analysis of germline stem cell proliferation. (D, F) Anterior portion of germaria from five-day-old control females showing germline stem cells in S phase (D, arrowhead) or M phase (F, arrowhead). EdU (D, magenta) labels cells in S phase and can be distinguished from fusome based on morphology and overlap with nucleus. Phospho-histone H3 (pHH3; F, green) labels cells in M phase of mitosis. (E, G) Frequencies of EdU-positive (E) or pHH3-positive (G) germline stem cells in five-day-old females on control or high sugar diets. Numbers of germline stem cells analyzed are shown inside bars. Data shown as mean±s.e.m. from three independent experiments. ns, no significant difference, Chi-square test. (H) Single confocal section of control ovariole containing β-gal-positive germline clones at three days after heat shock. Arrowhead indicates most developed labeled cyst (in a stage 7 follicle). β-gal (green) labels germline cysts originally labeled within anterior portion of germarium during heat shock (time zero; see Methods). Scale bar, 35 μm. (I) Box and whisker plot showing the stage of the most-developed germline cyst per ovariole at one and three days after heat shock in females on control or high sugar diets. At least fifty ovarioles were analyzed per sample for each time point. Data represent three independent experiments. ns, no significant difference, Chi-square test. (J, J’) Single confocal section showing a β-gal-positive follicle cell clone (arrowheads) at three days after heat shock. β-gal (J, green) labels follicle cells originated from single follicle cells labeled at time zero. (J’) β-gal channel shown in white. Scale bar, 50 μm. (K) Log scale plot showing the average number of follicle cells per clone over time. At least 85 follicle cell clones were analyzed per sample for each time point. Data shown as mean±s.e.m. from three independent experiments. ns, no significant difference, two-way ANOVA with interaction.

To determine if a high sugar diet affects the rate of follicle growth, we took advantage of a well-described lineage tracing system [22–25] (see Methods). We labeled dividing germ cells in the anterior region of germaria of females on control versus high sugar diets and followed their progress through oogenesis after one and three days (Fig 2H). The most developed labeled cysts reached similar stages in females on either diet, indicating similar growth and development rates (Fig 2I). We also measured the proliferation rates of follicle cells surrounding these growing germline cysts based on the increase in the size of clones generated from single labeled follicle cells and found no difference based on diet (Fig 2J and 2K). These results show that the growth of developing follicles is not affected by a high sugar diet.

### The ovaries of females on a high sugar diet have increased death of early germline cysts in germaria and degeneration of vitellogenic follicles

Having found that germline stem cell behavior and follicle growth are not altered on a high sugar diet (Fig 2), we asked whether high levels of dietary sugar might cause death of early germline cysts or follicles entering vitellogenesis. We labeled dying early germline cysts using the ApopTag TUNEL assay (Fig 3A) and observed a four-fold increase in the frequency of germaria with dying cysts in females maintained on high sugar versus control diets (Fig 3B). We identified degenerating vitellogenic follicles based on the presence of pyknotic nuclei (Fig 3C and D). Females on a high sugar diet had twice as many ovarioles containing degenerating vitellogenic follicles as females on a control diet (Fig 3E). Therefore, we conclude that the reduced egg production of females maintained on a high sugar diet is a direct consequence of the increased death of early germline cysts and vitellogenic follicles.

**Fig 3.**
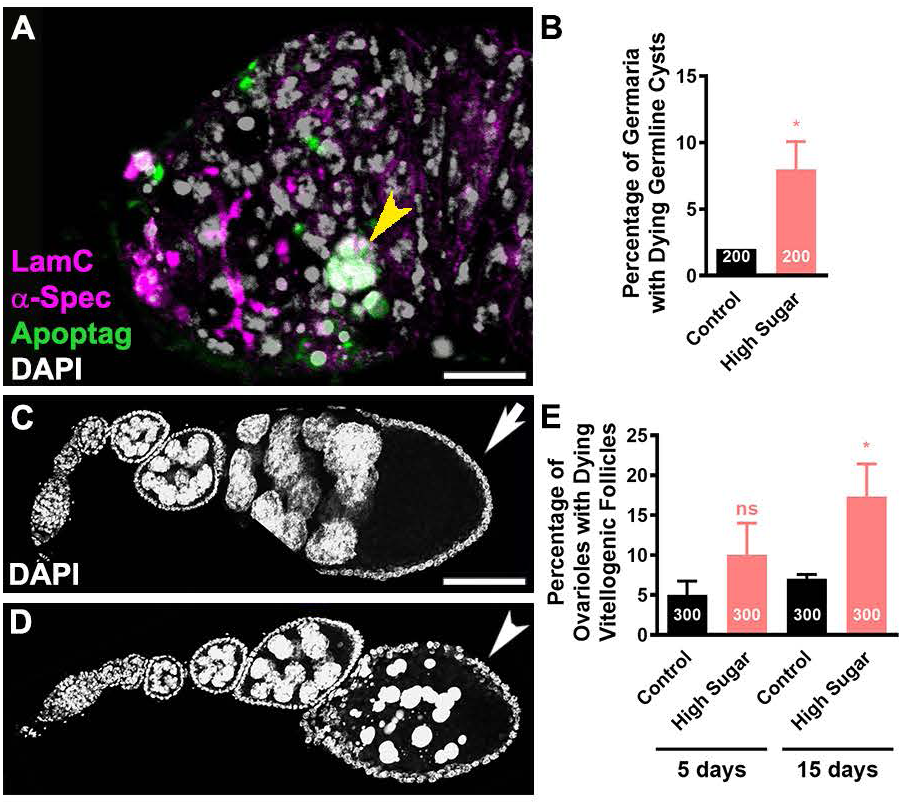
A high sugar diet increases death of early germline cysts and vitellogenic follicles. (A) Germarium from five-day-old female showing example of a dying early germline cyst (arrowhead). α-Spectrin (magenta) labels fusomes; Lamin C (magenta) labels cap cell nuclear envelopes; Apoptag (green) labels dying cells; DAPI (white) labels nuclei. Scale bar, 10 μm. (B) Frequencies of germaria containing Apoptag-positive dying cysts from five-day-old females on control or high sugar diets. Numbers of germaria analyzed are shown inside bars. Data shown as mean±s.e.m. from three independent experiments. \**P*<0.05, Chi-square test. (C, D) Ovarioles from 15-day old females showing healthy (C, arrow) or degenerating (D, arrowhead) vitellogenic follicles. Images represent projections of forty 1-μm confocal sections from 5×5 tiles. Scale bar, 25 μm. (E) Frequencies of ovarioles containing dying vitellogenic follicles from five- and 15-day-old females on control or high sugar diets. Numbers of ovarioles analyzed are shown inside bars. Data shown as mean±s.e.m. from three independent experiments. ns, no significant difference, \**P*<0.05, Chi-square test.

### Obesity is not sufficient to reduce female fertility

Many studies (including the present one) have shown that obesity induced by high dietary sugar leads to decreased fertility in *C. elegans*, *Drosophila*, and mammals [1,5,20,26] (Fig 1D-1I); however, it remains unclear whether these effects on fertility are a direct consequence of the diet, the obesity, or both. To determine if obesity alone is sufficient to impair fertility in *Drosophila* females, we generated obese females on a normal diet through genetic manipulation. We screened several genes involved in lipid metabolism (Table 1) using adult adipocyte-specific RNAi knockdown [27] and fat body triglyceride measurements [9]. Adult adipocyte-specific RNAi against *brummer* and *adipose* resulted in obese females (hereafter referred to as “genetically obese” females) (Fig 4A). Accordingly, the adipocytes of genetically obese females had much larger lipid droplets compared to control RNAi adipocytes (Fig 4B-E). Notably, *brummer* and *adipose* affect fat storage through distinct pathways: *brummer* encodes the homolog of mammalian ATGL and is the main *Drosophila* lipase for mobilization of lipids from lipid droplets [9], while *adipose* encodes the homolog of WDTC1, which binds to the DDB1-CUL4-ROC1 E3 ligase, targeting lipogenic enzymes for degradation [28]. We used both types of genetically obese females in our experiments to ensure that our results would be interpretable based on their obesity (as opposed to specific genes or pathways that are disrupted).

**Fig 4.**
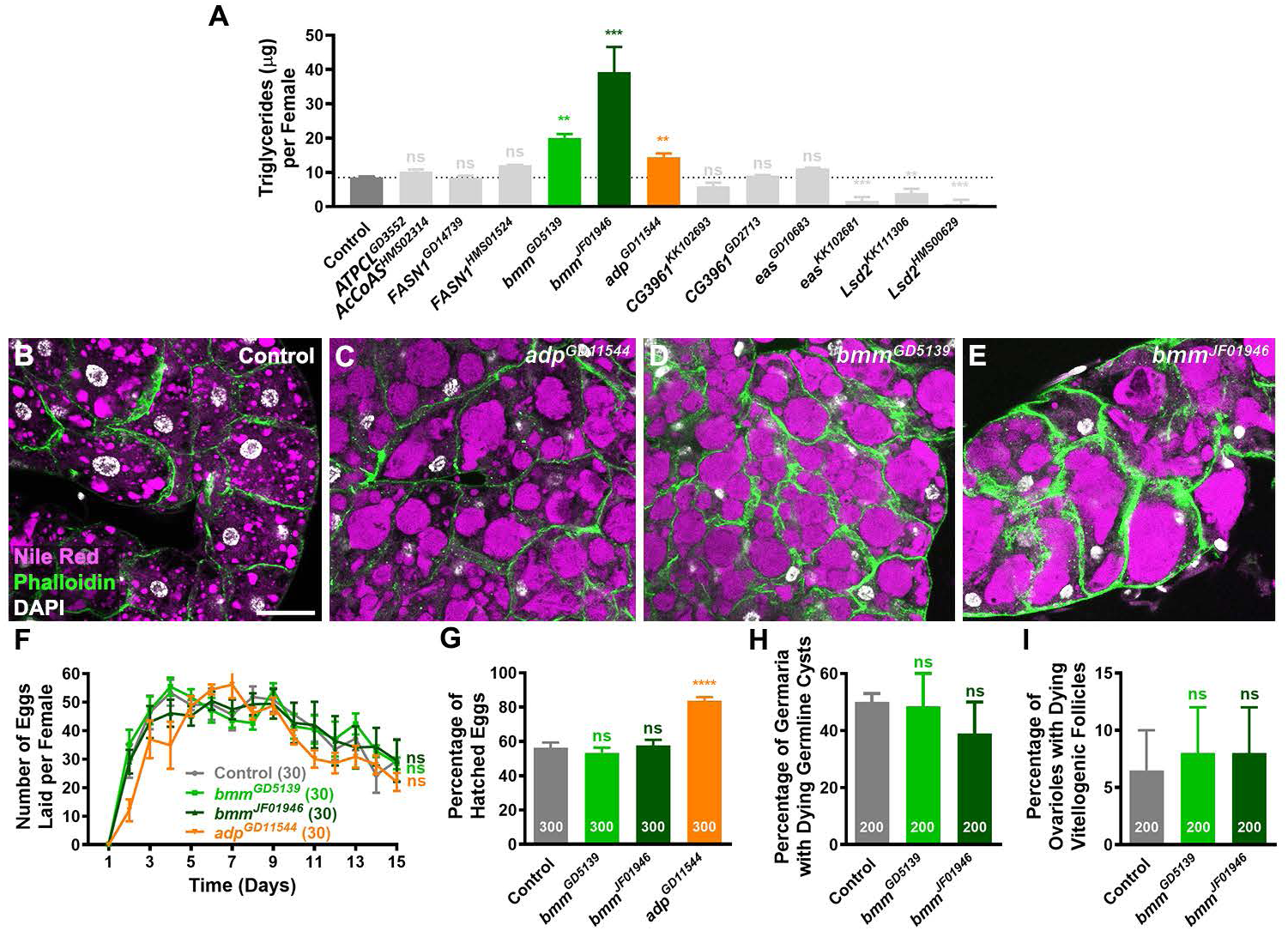
Obesity does not reduce female fertility. (A) Bar graph showing average triglyceride content per female (after subtraction of ovarian triglyceride content) of seven-day-old females with adult adipocyte-specific RNAi against control *LUC* or different genes involved in fat metabolism (see Table 1). Data shown as mean±s.e.m. from three independent experiments (10 females per genotype per experiment). \*\**P*<0.01; \*\*\**P*<0.001; ns, no significant difference, Student’s *t*-test. (B-E) Adipocytes from seven-day-old females with adult adipocyte-specific knockdown of *LUC* control (B), *adipose* (*adp*, C), or *brummer* (*bmm*, D, E). Nile red (magenta) labels lipid droplets; Phalloidin (green) outlines cell membranes; DAPI (white) labels nuclei. Images represent projections of three 1-μm confocal slices. Scale bar, 20 μm. (F) Line graph showing average number of eggs laid per female per day from one to 15 days of age. The number of females analyzed are shown in parentheses. Data shown as mean±s.e.m. from three experiments. ns, no significant difference, F-test of third order polynomial fitted curves. (G) Average percentage of eggs laid by females of different genotypes that hatch into larvae. Numbers of eggs analyzed are shown inside bars. Data shown as mean±s.e.m. from three experiments. \*\*\*\**P*<0.0001; ns, no significant difference, Student’s *t*-test. (H) Frequencies of germaria containing Apoptag-positive cysts from five-day-old females of different genotypes. Numbers of germaria analyzed are shown inside bars. Data shown as mean±s.e.m. from three independent experiments. ns, no significant difference, Chi-square test. (I) Frequencies of ovarioles containing dying vitellogenic follicles from 15-day-old females of different genotypes. Numbers of ovarioles analyzed are shown inside the bars. Data shown as mean±s.e.m. from three independent experiments. ns, no significant difference, Chi-square test.

**Table 1.** *UAS* lines used for RNA interference in this study.

| *UAS* line | Genotype | Source | Reference |
| --- | --- | --- | --- |
| *Luciferase*<sup>JF01355</sup><br>(*LUC*<sup>JF01355</sup><br>control) | *y<sup>1</sup> v<sup>1</sup>; P{TRiP.JF01355}attP2* | BDSC#31603 | [40,58] |
| *ATPCL*<sup>GD3552</sup> | *y<sup>1</sup> w<sup>1118</sup>; P{GD3552}v30282* | VDRC#30282 | [40,59] |
| *AcCoAS*<sup>HMS02314</sup> | *P{TRiP.HMS02314}/TM3* | BDSC#41917 |  |
| *FASN1*<sup>GD14739</sup> | *w<sup>1118</sup>; P{GD14739}v29349/CyO* | VDRC#29349 | [60] |
| *FASN1*<sup>HMS01524</sup> | *y1 sc\* v1; P{TRiP.HMS01524}attP2* | BDSC#35775 | [61] |
| *bmm*<sup>GD5139</sup> | *w<sup>1118</sup>; P{GD5139}v37877* | VDRC#37877 | [62] |
| *bmm*<sup>JF01946</sup> | *y<sup>1</sup> v<sup>1</sup>; P{TRiP.JF01946}attP2/TM3, Sb1* | BDSC#325926 | [58,63] |
| *adp*<sup>GD11544</sup> | *w<sup>1118</sup> P{GD11544}v22008* | VDRC#22008 | [64] |
| *CG3961<sup>KK102693</sup>* | *P{KK102693}VIE-260B* | VDRC#107281 | [40,64] |
| *CG3961<sup>GD2713</sup>* | *w<sup>1118</sup>; P{GD2713}v37305* | VDRC#37305 | [40,64] |
| *eas<sup>GD10683</sup>* | *w<sup>1118</sup>; P{GD10683}v34287/TM3* | VDRC#34287 | [40,64] |
| *eas<sup>KK102681</sup>* | *P{KK102681}VIE-260B* | VDRC#103784 | [40,64] |
| *Lsd2<sup>KK111306</sup>* | *P{KK111306}VIE-260B* | VDRC#102269 | [64,65] |
| *Lsd2<sup>HMS00629</sup>* | *y<sup>1</sup> sc<sup>\*</sup> v<sup>1</sup>; P{TRiP.HMS00629}attP2* | BDSC#32846 | [64,65] |

We measured fertility and examined specific stages and processes of oogenesis in genetically obese females as described above. Genetically obese females did not display any reduction in egg production or hatching rates relative to control RNAi females (Fig 4F and 4G). Accordingly, germline stem cell maintenance and proliferation (Fig S1), early germline cyst death (Fig 4H), and follicle degeneration (Fig 4I) were similar in control and genetically obese females. These results demonstrate that obesity alone does not negatively impact oogenesis.

### Obesity induced by high dietary sugar is resistant to leanness-inducing genetic manipulations

To determine if a high sugar diet is sufficient to reduce fertility or if it requires co-occurring obesity, we would ideally maintain females on a high sugar diet while preventing them from becoming obese. We attempted to genetically suppress obesity in females maintained on a high sugar diet by overexpressing the lipase-encoding gene *brummer* while simultaneously inducing RNAi against *Lsd2* in adult adipocytes (“high *brummer* low *Lsd2*”). Lsd2 encodes the homolog of mammalian perilipin, a protein that envelops lipid droplets and limits Brummer access [29]. Therefore, “high *brummer* low *Lsd2*” represents a powerful genetic manipulation for inducing leanness. (Also see Fig 4A for effect of *Lsd2* knockdown alone on adiposity of females on a control diet.) Remarkably, the levels of adipocyte lipid storage in “high *brummer* low *Lsd2*” females were just as high as in control females on a high sugar diet (Fig S2). These results underscore the challenges of reversing high sugar-induced obesity and precluded our intended analysis of non-obese females on a high sugar diet.

### Glucose, trehalose, and glycogen levels are increased in high-sugar-obese but not genetically obese females

Females maintained on a high sugar diet and adipocyte *brummer* knockdown females on a normal diet were similarly obese (Fig 1D-1F and Fig 4A-4E), but only high sugar females had impaired oogenesis (Fig 1H and 1I and Fig 4F and 4G). Previous studies showed that glycogen and trehalose levels are increased in high-sugar-obese females [14,20] and that the levels of glycogen do not change in *brummer^1^* global mutants [9]. Therefore, to investigate possible reasons for the difference in fertility between females on high sugar versus adipocyte *brummer* knockdown females, we compared their levels of glycogen, trehalose and glucose relative to their respective controls. In agreement with the literature, we observed that glycogen levels were doubled in high-sugar-obese females compared to their controls (Fig 5A) but remained unchanged in genetically obese females compared to control RNAi females (Fig 5B). In addition, we also found that trehalose levels in high-sugar-obese females were four-fold those of their controls (Fig 5C), while trehalose was unaffected in genetically obese females relative control RNAi females (Fig 5D). Moreover, our results show a three-fold increase in glucose levels in high-sugar-obese females (Fig 5E) but no consistent change in genetically obese females (Fig 5F). Thus, high-sugar-obese females, but not genetically obese females, had increased levels of glycogen, trehalose, and glucose, showing a correlation between high carbohydrate levels in females and reduced fertility.

**Fig 5.**
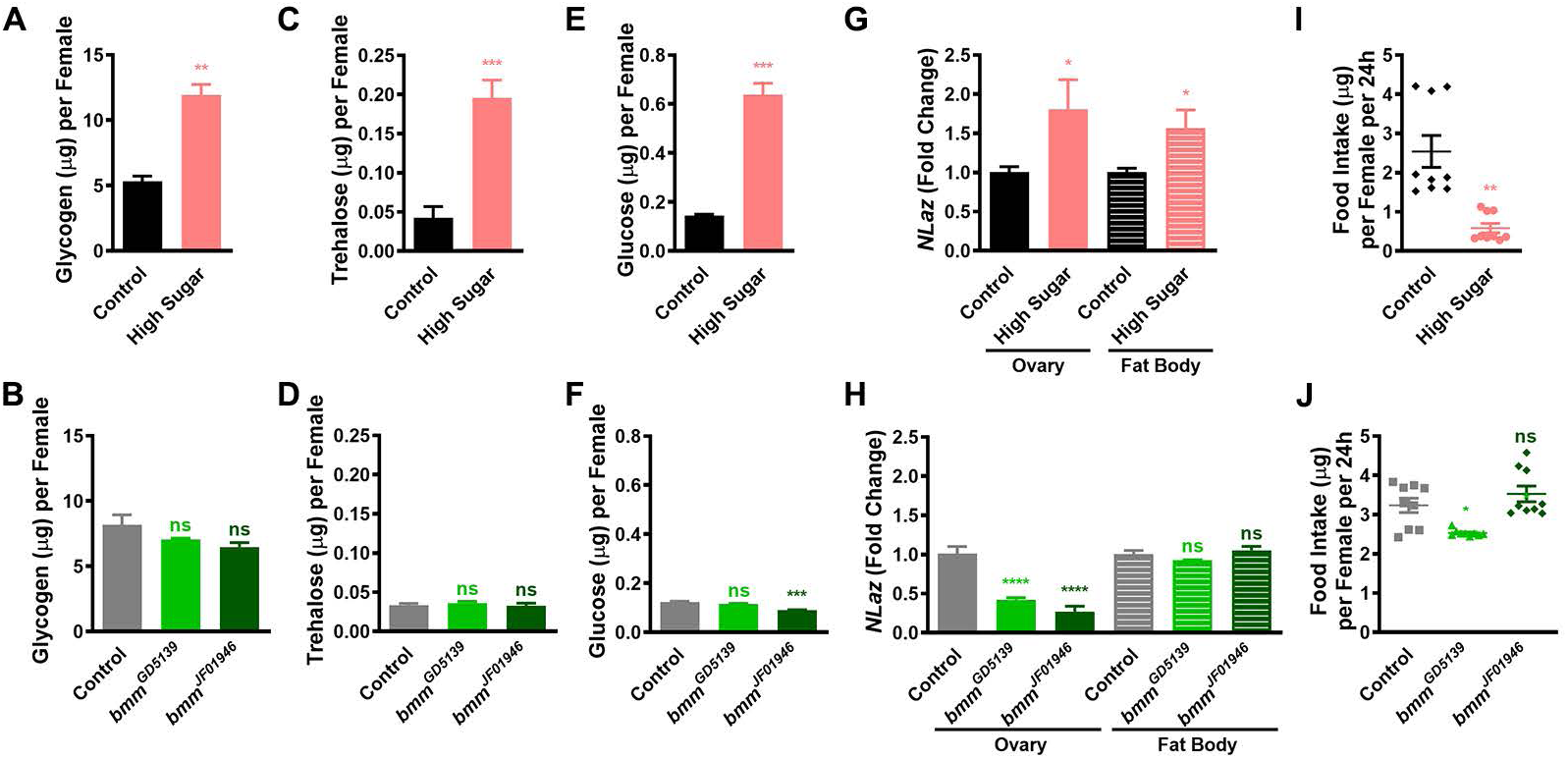
High sugar diet, but not genetically induced obesity, leads to increased glucose, trehalose, and glycogen levels, insulin resistance, and reduced food consumption. (A-F) Average glycogen (A, B), trehalose (C, D), and glucose (E, F) content of seven-day-old females on different diets (A, C, E) or of different genotypes (B, D, F). Data shown as mean±s.e.m. from three experiments (10 females per sample per experiment). \*\*\**P*<0.001; \*\**P*<0.01; ns, no significant difference, Student’s *t*-test. (G, H) Bar graphs showing relative levels of the insulin resistance marker *NLaz* (based on quantitative real-time PCR) in ovaries or fat bodies of seven-day-old females on different diets (G) or of different genotypes (H). Data shown as mean±s.e.m. from three experiments (10 females per sample per experiment). \*\*\*\**P*<0.0001; \**P*<0.05, Student’s *t*-test. (I, J) Dot plots showing food consumption per female per 24 hours for seven-day-old females on different diets (I) or of different genotypes (J). Each dot represents the average consumption from 10 females. The horizontal line represents mean±s.e.m. from three experiments with three biological replicates each (see methods). \*\**P*<0.01; ns, no significant difference, Student’s *t*-test.

### Insulin resistance in the fat body and ovary is increased in high-sugar-obese but not genetically obese females

Sustained increases in circulating sugars are typically associated with insulin resistance (i.e., an impaired response to insulin) in adipocytes in *Drosophila* and mammals [30]. We previously showed that insulin signaling in adipocytes and/or ovaries promotes the survival of early germline cysts and vitellogenic follicles, among its other roles [18,19,31]. We therefore asked if the high trehalose and glucose levels we observed in high-sugar-obese females (Fig 5C-5F) lead to insulin resistance in the fat body and/or ovary. The Lipocalin-encoding gene *Neural Lazarillo* (*NLaz*), which is required for high sugar diet-induced insulin resistance in larvae [32], is routinely used as a marker of insulin resistance in *Drosophila* larvae and adults [32–35]. We found that *NLaz* mRNA levels were significantly higher in the fat body and ovary of high-sugar-obese females (Fig 5G) but were unchanged in the fat body and reduced in the ovaries of genetically obese females (Fig 5H) compared to their respective controls. These correlative results raise the formal possibility (albeit later disproved; see below) that reduced insulin signaling resulting from increased insulin resistance might contribute to the increased death of early germline cysts and vitellogenic follicles observed in high-sugar-obese females.

### High-sugar-obese females consume less overall food but more sugar than controls

To address whether differences in food consumption might also contribute to the metabolic and ovarian phenotypes of high-sugar-obese females, we measured their food intake using the Consumption-Excretion method [23,36]. High-sugar-obese females ate one-quarter as much as their controls on a normal diet (Fig 5I), in agreement with previous studies [20]. However, given that the concentration of sucrose in the high sugar food is six-fold that of the control food, females on the high sugar diet ingested higher levels of sugar compared to their controls. By contrast, genetically obese females showed similar food intake to control RNAi females (Fig 5J). These results suggest that, in principle, increased sugar consumption, reduced intake of other nutrients, or both might potentially contribute to the reduced fertility of high-sugar-obese females.

### Hydration restores normal fertility and glucose levels in high-sugar-obese females independent of food intake, glycogen levels, insulin resistance, or obesity

A recent study showed that providing an additional source of dietary water reverses the negative effects of a high sugar diet on the lifespan of *Drosophila* females, although these females remain obese with high levels of trehalose and insulin resistance [20]. We therefore asked if supplementation of high sugar diet with an *ad libitum* water source might also reverse the fertility defects of high-sugar-obese females. We first examined the effects of dietary water supplementation on the metabolic parameters of females on a high sugar diet. As previously reported [20], triglyceride levels were equally high in females on a high sugar diet with or without water supplementation (Fig 6A). Interestingly, glycogen levels also remained high in females on a high sugar diet regardless of extra dietary water (Fig 6B). By contrast, water supplementation reduced the levels of trehalose by 17% (Fig 6C) and glucose by 58% (Fig 6D) in females on a high sugar diet. Remarkably, the levels of insulin resistance based on the *NLaz* marker nearly doubled in the ovaries and were slightly elevated in the fat bodies of females on a high sugar diet supplemented with water compared to those without water supplementation (Fig 6E). As a second approach, we measured the levels of phosphorylated AKT kinase (pAKT), the active form of a downstream component of the insulin pathway, which is commonly used to monitor insulin resistance [20,37]. In both the ovaries and fat bodies of females on a high sugar diet, the increase (or lack thereof) in pAKT levels in response to added insulin was statistically similar regardless of water supplementation (Fig 6F-6G’), similar to a previous report for fat bodies [20]. Despite discrepancies in the specific results using *NLaz* versus pAKT to assess insulin resistance, both approaches clearly showed that water supplementation does not reverse the high sugar diet-induced insulin resistance in either fat bodies or ovaries. Food intake on a high sugar diet was not altered by water supplementation (Fig 6H), as previously reported [20]. Altogether, our analysis indicates that dietary water supplementation drastically reverses the high glucose levels (with a smaller effect on trehalose) but not the insulin resistance of females on a high sugar diet, with no significant effects on their glycogen levels, food intake, or obesity.

**Fig 6.**
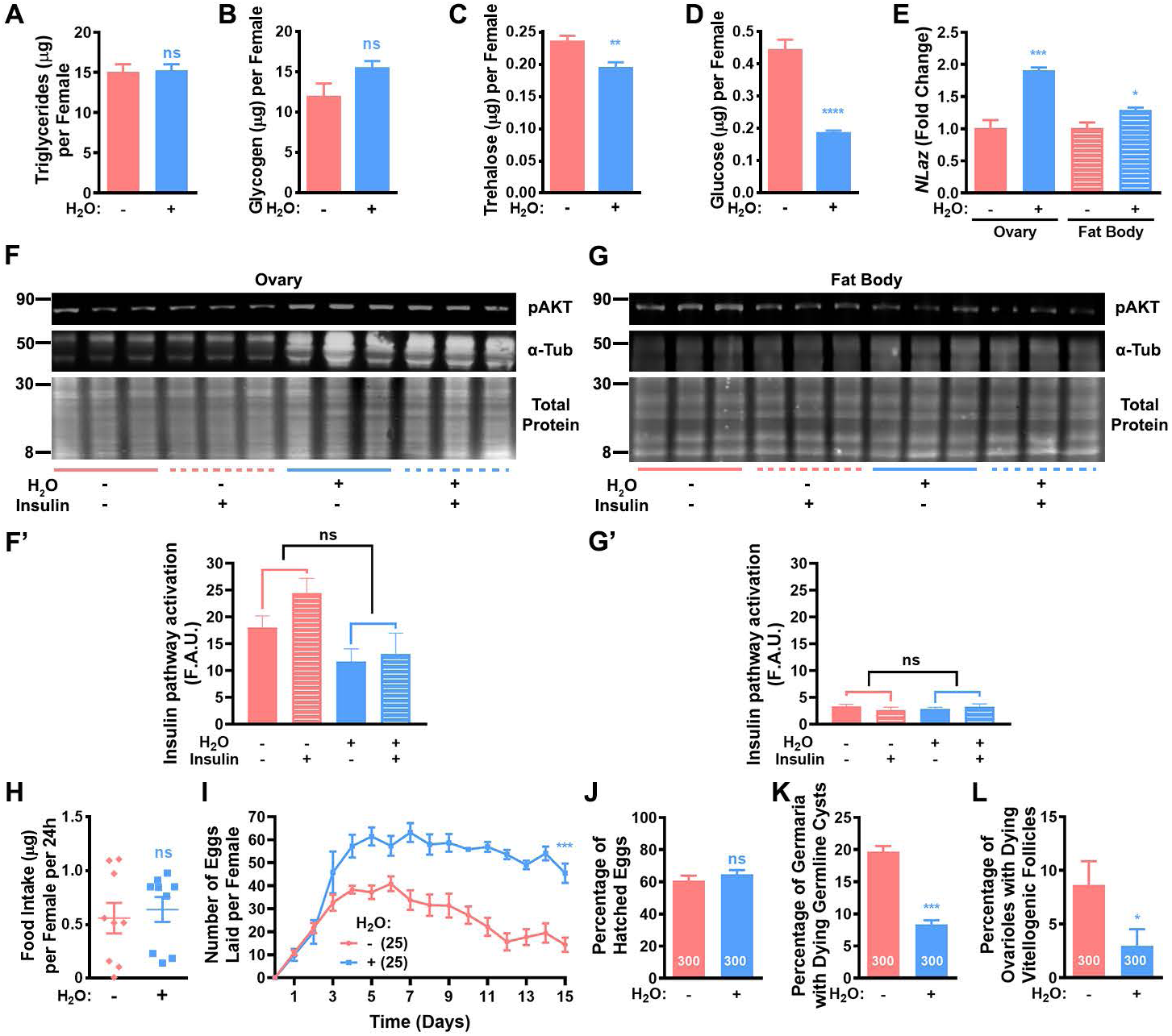
Dietary water supplementation restores normal glucose levels and fertility of females on a high sugar diet independent of food consumption, insulin resistance, or obesity. (A-D) Average triglyceride (after subtraction of ovarian triglyceride; A), glycogen (B), trehalose (C), and glucose (D) content of seven-day-old females on a high sugar diet with (+) or without (-) extra dietary water (H_2_O). Data shown as mean±s.e.m. from three experiments (10 females per sample per experiment). \*\*\*\**P*<0.0001; \*\**P*<0.01; ns, no significant difference, Student’s *t*-test. (E) Relative levels of the insulin resistance marker *NLaz* (based on quantitative real-time PCR) in ovaries or fat bodies of seven-day-old females on a high sugar diet with or without dietary water supplementation. Data shown as mean±s.e.m. from three experiments (10 females per sample per experiment). \*\*\**P*<0.001; \**P*<0.05, Student’s *t*-test. (F,G) Western blots of extracts from ovaries (F) or fat bodies (G) [incubated with (+) or without (-) insulin for 20 minutes; see Methods] from seven-day old females on a high sugar diet with or without water supplementation. Phosphorylated AKT kinase (pAKT) shown in top panels; *α*-Tubulin (*α*-Tub) shown in middle panels; total protein shown in bottom panels. (F’, G’) Bar graphs showing average densitometry of pAKT normalized for total protein for blots in (F, G), respectively. *α*-Tub was not used for normalization owing to its higher expression with extra dietary water. Data shown as mean±s.e.m. from three experiments. Fold-changes in response to insulin were statistically compared. ns, no significant difference, two-way ANOVA with interaction. (H) Dot plot showing food consumption per female per 24 hours for seven-day-old females on a high sugar diet with or without water supplementation. Each dot represents the average consumption from 10 females. The horizontal line represents mean±s.e.m. from three experiments with three biological replicates each. ns, no significant difference, Student’s *t*-test. (I) Average number of eggs laid daily per female on a high sugar diet with or without water supplementation. The numbers of females (paired with males) analyzed are shown in parentheses. Data shown as mean±s.e.m. from three experiments. \*\*\**P*<0.001, *F*-test of third order polynomial fitted curves. (J) Average percentage of eggs laid by females on a high sugar diet with or without dietary water supplementation that hatch into larvae. The number of eggs analyzed are shown inside bars. Data shown as mean±s.e.m. from three experiments. ns, no significant difference, Student’s *t*-test. (K) Frequencies of germaria containing Apoptag-positive dying cysts from five-day-old females on a high sugar diet with or without water supplementation. Numbers of germaria analyzed are shown inside bars. Data shown as mean±s.e.m. from three independent experiments. \*\*\**P*<0.001, Chi-square test. (L) Frequencies of ovarioles containing dying vitellogenic follicles from 15-day-old females on a high sugar diet with or without water supplementation. Numbers of ovarioles analyzed are shown inside the bars. Data shown as mean±s.e.m. from three independent experiments. \**P*<0.05, Chi-square test.

Next, we compared the fertility of high-sugar-obese females with or without dietary water supplementation. Interestingly, extra dietary water significantly increased egg production of females on a high sugar diet (Fig 6I), while hatching rates remained similar regardless of water supplementation (Fig 6J). These results indicate that a high sugar diet impairs egg production and hatching rates through different mechanisms. In accordance with the increase in egg production, water supplementation significantly decreased the frequency of dying early germline cysts and vitellogenic follicles in high-sugar-obese females (Fig 6K and 6L).

Taken together, these findings lead to major conclusions regarding the potential mechanisms mediating the negative effects of a high sugar diet on oogenesis. First, there is a strong correlation between high glucose levels and lower egg production, suggesting a possible causal relationship. Second, despite their reduced food consumption and elevated insulin resistance, high-sugar-obese females have sufficient nutrient intake and insulin signaling to support oogenesis. Third, females on a high sugar diet supplemented with water remain obese yet have largely restored egg production, strengthening our conclusion that obesity does not directly cause the observed oogenesis defects – namely, germline cyst death and follicle degeneration – observed in high-sugar-obese females.

## DISCUSSION

From *Drosophila* to humans, many studies have established a strong association between obesity and reduced fertility [1,5,6,26,38,39]. However, the methods for generating obese animals in these studies introduce major confounding factors, such as diets high in sugar and/or fat or hormonal changes leading to increased appetite. For example, a high sugar diet causes obesity in *Drosophila*, and these obese females produce fewer eggs [5,6], making it unclear if a high sugar diet, obesity, or both reduce their fertility. In this study, we show that high-sugar-obese *Drosophila* females produce fewer eggs because of increased death of early germline cysts and degeneration of vitellogenic follicles (Fig 7). Further, we demonstrate that this reduction in fertility is not caused by obesity, insulin resistance, or high trehalose or glycogen levels (Fig 7). Instead, we find a strong correlation between high glucose levels and the decreased egg production of females on a high sugar diet, as supplementation with dietary water drastically improves both glucose levels and egg production, with much smaller or no effects on other metabolic parameters (Fig 7). Our findings provide a foundation for future studies to investigate the causal relationship between high glucose levels (and/or other dietary water-dependent factors) and reduced *Drosophila* fertility. More broadly, they also highlight the importance of carefully controlling for confounding factors when investigating how obesity affects the risk of infertility and other obesity-associated disorders, including chronic inflammation, cardiovascular abnormalities, and cancers [30,37].

**Fig 7.**
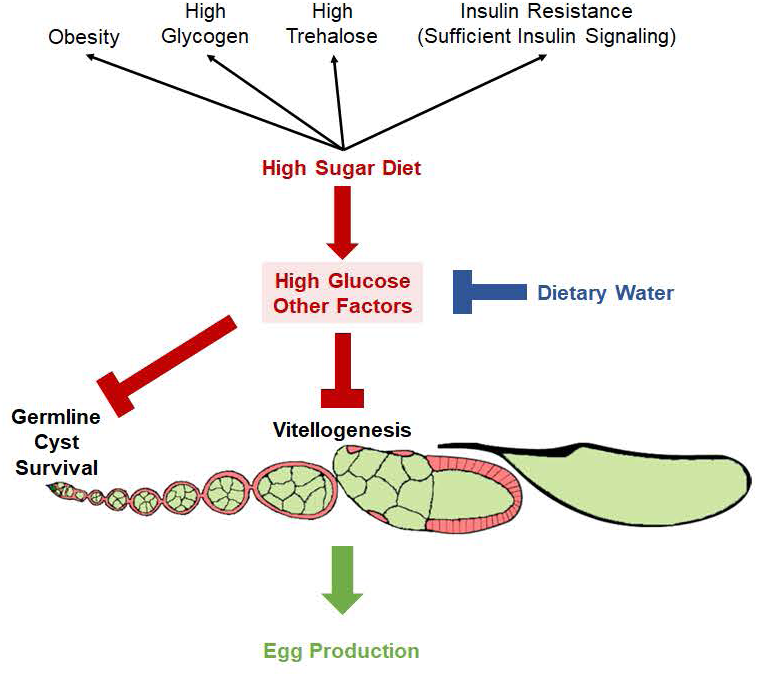
Model for how a high sugar diet reduces *Drosophila* female fertility. A high sugar diet leads to increased death of early germline cysts and vitellogenic follicles, thereby reducing egg production. The negative effects of a high sugar diet on oogenesis are mediated by elevated glucose and/or changes in other factors that are reversible by dietary water supplementation. Conversely, elevated adiposity, glycogen, trehalose or insulin resistance do not correlate with lower egg production on a high sugar diet.

### Excess fat accumulation in adipocytes does not disrupt their function in *Drosophila*

Our previous published studies have shown that adipocytes are integral to the physiological regulation of *Drosophila* oogenesis [16]. For example, dietary yeast rapidly changes the expression of enzymes in multiple metabolic pathways in adipocytes, and adipocyte-specific knockdown of key components of these pathways leads to specific phenotypes, such as germline stem cell loss, early cyst death, or degeneration of vitellogenic stages [40]. Reduced amino acid sensing by adipocytes reduces germline stem cell numbers and inhibits ovulation [27], while blocking insulin signaling specifically in adipocytes increases loss of germline stem cells, death of early germline cysts and degeneration of vitellogenic follicles [18]. Our current findings indicate that none of these known roles of adipocytes are impaired despite very high levels of fat accumulation in genetically obese females. We conclude that obesity in and of itself does not impair adipocyte function.

This work underscores the importance of designing studies that tease apart the contributions of obesity itself from obesity-independent effects of obesogenic diets. Between 1980 and 2018, the percentage of obese American adults rose from 19% to 42% for women, and from 13% to 43% in men [41]. This rapid rise in obesity was largely a consequence of unhealthy diets and/or lack of physical exercise [42] (www.cdc.gov). Therefore, dietary and/or physical activity factors are likely contributing to the adverse health outcomes of a large fraction of adults that are currently obese. Meanwhile, ∼10 to 30% of obese individuals are clinically recognized as having metabolically healthy obesity (i.e. obesity with normal glucose and lipid metabolism parameters and absence of hypertension) for unclear reasons [43]. Yet, most studies in human populations or laboratory animal models examining the connection between obesity and infertility do not clearly distinguish adiposity and dietary contributions to their findings [1,5,6,14,26,32,34,44–48]. For example, to our knowledge, the effect of obesity caused by mutation of *Wdtc1* [11] or *Atgl* [48] on the fertility of mammalian females has not been studied. It would be interesting to compare findings from such types of studies to our findings of normal fertility in genetically obese females.

### Insulin resistance markers do not necessarily indicate insufficient levels of insulin signaling

Insulin signaling is crucial for the regulation of *Drosophila* oogenesis. In addition to the indirect roles of insulin signaling within adipocytes to control ovarian processes noted above, insulin-like peptides also act directly on the germline to control germline stem cell proliferation, follicle growth and vitellogenesis [19,49], and on cap cells to control germline stem cell maintenance [17,49,50]. In this study, we showed that insulin resistance markers are elevated in the fat body, as previously described [32,34], and in the ovaries of females on a high sugar diet; yet, only germline cyst and vitellogenic follicle survival are affected in these females. Based on our previous genetic mosaic analyses, we can rule out the possibility that these two processes have higher threshold requirements for insulin signaling than other insulin-dependent processes. Specifically, germ cells that are homozygous mutant for the hypomorphic insulin receptor allele *InR^353^* allow survival of most vitellogenic follicles (in contrast to the death of all follicles containing *InR^E19^* or *InR^339^* mutant germ cells), while germline stem cell maintenance and follicle growth remained significantly impaired [19]. Further, dietary water supplementation of a high sugar diet led to increased levels of insulin resistance markers in both the ovary and fat body, while reversing the death phenotypes of early cysts and vitellogenic follicles. Altogether, we can definitively conclude that females on a high sugar diet have sufficient levels of insulin signaling in both their ovaries and adipocytes to support normal rates of oogenesis. These findings raise the possibility that in other systems where insulin resistance is observed [1,6,20,32,34,48,51], there might be sufficient levels of insulin signaling for normal functioning of the tissues in question.

### High sugar levels and dehydration

Biological water participates in various processes, including metabolism [52–54], and a correlation between obesity and dehydration has been reported in humans [55], although it remains unclear whether dehydration correlates with obesity itself or with obesogenic diets. Notably, in *Drosophila*, dietary water supplementation reverses the shortened lifespan and dehydration (i.e., decreased hemolymph volume) caused by a high sugar diet, even though flies remain obese and have high levels of trehalose and insulin resistance [20]. Here, we further show that dietary water supplementation drastically reversed the high glucose levels and reduced fertility of *Drosophila* females on a high sugar diet. Future studies should address whether high glucose levels have a causal relationship with reduced fertility and, if so, whether the ovaries sense glucose levels, decreased hemolymph volume, or other signals.

## MATERIALS AND METHODS

### *Drosophila* strains and experimental conditions

*Drosophila* stocks were maintained at 22°C on standard medium composed of 4.64% w/v yellow cornmeal (Quaker), 5.8% v/v unsulphured cane molasses (Sweet Harvest Feeds),1.74% w/v active dry yeast (<u>Red Star</u>) and 0.93% w/v agar (Apex BioResearch Products). (Note: The concentration of 5.8% molasses in our standard medium corresponds to ∼5% sugar, mostly sucrose.) The *y w* “wild-type” strain was used for all experiments, except where indicated. The adipocyte-specific *3.1Lsp2-Gal4* [56] and *tub-Gal80^ts^* [57] were used in combination to induce adult-specific RNAi as described [27] using the *UAS-RNA hairpin* lines listed on Table 1. The *X-15-*29, X*-15-33*, and *MKRS, hs-FLP* lines used for lineage tracing experiments have been described [24]. For most experiments, flies were raised at 22°C on standard medium and zero-to-one-day-old adults were incubated at 25°C, ≥70% humidity, on standard medium (∼5% sugar content; control diet) or standard medium with added sucrose (∼32% sugar content; high sugar diet). For adult adipocyte-specific experiments, flies were raised at 18°C on standard medium and zero-to-one-day-old adults were incubated at 29°C, ≥70% humidity, on standard medium. For dietary water supplementation, a barrier tip (20 µl; Genesse Scientific) filled with 1% agar was inserted in the food and taped to the wall of the vials containing high sugar medium. For all experiments, fly medium was supplemented with dry yeast and changed daily (along with the 1% agar-filled tip, if applicable).

### Food consumption assay

To measure food intake, we used the consumption-excretion dye-based method [36] with minor modifications, as described previously [23]. Briefly, 10 zero-to-one-day-old couples were maintained on control or high sugar medium for six days at 25°C (or 29°C, for RNAi experiments). Females were then transferred to empty vials closed with plugs containing (on their internal surface) ∼0.5 ml control or high sugar medium with 0.25% FD&C Blue No. 1 (Spectrum Chemicals), respectively, for 24 hours. (A similar procedure was followed to control for background absorbance, except that media on plugs did not have the blue dye.) To recover the internal dye, females were homogenized in 1.5 ml water, homogenates were centrifuged at 10,000 g for 1 minute, and supernatants were collected. To collect the dye excreted by females, 3 ml water were used to rinse the vials and water extracts were vortexed for 10 seconds. Absorbance at 630 nm was measured using a Synergy H1 spectrophotometer (Agilent BioTek) and converted to amount of medium consumed (µg/fly) based on a standard curve of pure FD&C Blue No. 1 in water. Experiments were done in triplicate and statistical analysis was performed using the Student’s *t*-test (GraphPad Prism).

### Quantification of laid eggs and egg hatching

To count the number of eggs laid, five couples were maintained in inverted perforated plastic bottles closed with control or high sugar medium plates supplemented with dry yeast, in six replicates, at 25°C (or 29°C, for RNAi experiments). Plates were changed twice a day, and eggs laid within a 24-hour period were counted daily for 15 days to calculate the average number of eggs produced per female per day. Experiments were done in triplicate and statistical analysis was performed using F-test of third order polynomial (GraphPad Prism) fitted curves.

To quantify hatching rates, eggs laid overnight from seven-day to eight-day time points of the egg count experiments above were collected. For each experiment, 10 groups of 10 eggs per condition were placed on a molasses plate around a dab of yeast paste and incubated at 25°C for 24 hours in a humid chamber in triplicate. The unhatched eggs were counted and subtracted from the total to calculate the number of hatched eggs. Experiments were done in triplicate and statistical analysis was performed using the Student’s *t*-test (GraphPad Prism).

We note that our measurements of egg laying and hatching rates fall within a higher range than previously reported [5]. These discrepancies are likely due to differences in methodology, such as density of flies and/or food composition.

### Tissue staining and microscopy

Ovaries were dissected and ovarioles were teased apart in Grace’s Insect Medium (Bio Whittaker). Samples were fixed for 15 minutes at room temperature in fixing solution [5.3% paraformaldehyde (Ted Pella) in Grace’s medium], and then rinsed and washed three times for 15 minutes each in PBST [0.1% Triton X-100 in PBS (10 mM NaH_2_PO_4_/NaHPO_4_, 175 mM NaCl, pH 7.4)]. Samples were blocked for three hours at room temperature in blocking solution [5% normal goat serum (MP Biochemicals) plus 5% bovine serum albumin (Sigma) in PBST] and then incubated overnight at 4°C in primary antibodies diluted in blocking solution: 1:20 mouse monoclonal anti-α-Spectrin [3A9, Developmental Studies Hybridoma Bank (DSHB)]; 1:20 mouse monoclonal anti-Lamin C (LC28.26, DSHB); 1:200 chicken anti-β-gal antibody (ab9361, Abcam); or 1:200 rabbit anti-phospho-histone H3 (Ser10) (06-570, Millipore). Ovaries were washed as described and incubated with 1:400 secondary antibodies conjugated to Alexa Fluor 488 or Alexa Fluor 568 (A11034 or A11004, respectively, Molecular Probes – Invitrogen) in blocking solution for two hours (protected from light) at room temperature. Samples were washed (protected from light) and mounted in Vectashield containing DAPI (4’,6-diamidino-2-phenylindole, a fluorescent stain specific for DNA) (Vector Labs). For lipid droplet visualization, abdominal carcasses (without ovaries or guts) dissected in Grace’s medium were fixed for 20 minutes at room temperature in fixing solution, and then rinsed and washed three times for 15 minutes each in PBST. Carcasses were then incubated with 1:200 Alexa fluor 488-conjugated phalloidin (a bicyclic peptide that binds F-actin) (A12379, Invitrogen) for 40 minutes at 4°C and washed as described above. Samples were then stained with 25 ng/mL Nile Red (a lipophilic fluorescent dye that stains lipid droplets) (19123, Sigma) in Vectashield containing DAPI for at least 48 hours at 4°C. All experiments were done in triplicate, and data were collected using a Zeiss AxioImager-A2 fluorescence microscope or Zeiss LSM700 or LSM900 confocal microscopes.

### Quantification of germline stem cell and cap cell numbers and germline stem cell proliferation

Cap cells were identified based on their ovoid nuclei and Lamin C-positive staining, while germline stem cells were identified based on their juxtaposition to cap cells and typical fusome morphology and position [66]. Experiments were done in triplicate and statistical analysis was performed using two-way ANOVA with interaction (GraphPad Prism).

To label germline stem cells in S phase, we used an EdU (5-ethynyl-2’-deoxyuridine; a nucleoside analog of thymidine) incorporation assay. Intact dissected ovaries were incubated for one hour at room temperature in 100 μM EdU (Molecular Probes) in Grace’s insect medium, washed, and fixed as described above. Following incubation with primary antibodies (anti-phospho-histone H3, anti-Lamin C, and anti-α-Spectrin), samples were subjected to the Click-iT reaction according to the manufacturer’s instructions (Life Technologies) for 30 minutes at room temperature. Ovaries were washed, incubated with secondary antibodies, and washed again prior to mounting and microscopy, as described above. We calculated the fraction of EdU-positive or phospho-histone H3-positive germline stem cells as a percentage of the total number of germline stem cells analyzed per condition. Experiments were done in triplicate and statistical analysis was performed using Chi-square analysis.

### Assessment of follicle growth and development using lineage tracing analysis

Follicle development through oogenesis involves the growth of the germline cysts (originally produced by four incomplete mitotic divisions of cystoblasts in the anterior portion of the germarium) in coordination with mitotic divisions of surrounding follicle cells (until stage 7 of oogenesis, when follicle cells transition to endoreplication) [67]. β-galactosidase (β-gal)-positive clones from single mitotically dividing cells were produced as previously described [22,23]. Newly eclosed *y w; X-15-29/X-15-33; MKRS, hs-FLP*/+ females (with zero-to-one-day-old *y w* males) were maintained on control or high sugar diets for two days at 25°C. Flies were then heat-shocked at 37°C for one hour to induce flippase expression (and the generation of single β-gal-positive cells at day zero) and subsequently transferred to their respective media (changed daily) for one or three days prior to ovary dissections. To assess follicle growth, we analyzed β-gal-positive germline and follicle cell clones at both time points. For germline clones (i.e. partially or fully labeled 16-cell cysts generated from mitotically dividing germline stem cells, cystoblasts, or two-, four-, or eight-cysts present at day zero), we identified the most developed follicle stage containing a partially/fully β-gal labeled cyst in each ovariole analyzed. Statistical analysis was performed using Chi-square analysis. For follicle cells (i.e. β-gal-positive clones generated from single follicle cells labeled at day zero), the number of labeled cells per clone was counted in stages 4 to 6 follicles. Doubling times were calculated using regression line equations (GraphPad Prism). Experiments were done in triplicate and statistical analysis was performed using two-way ANOVA with interaction (GraphPad Prism).

### Analysis of early dying cysts and degenerating vitellogenic follicles

To quantify germaria containing dying germline cysts, we used the ApopTag Fluorescein Direct In Situ Apoptosis Detection Kit (S7160, Millipore Sigma) as previously described [22]. Degenerating vitellogenic follicles were identified in DAPI-stained ovarioles based on the presence of pyknotic nuclei, which are not present in healthy vitellogenic follicles. Experiments were done in triplicate and statistical analysis was performed using Chi-square analysis.

### Triacylglycerol measurements

For triacylglycerol (TAG) quantification, 10 whole females or 10 pairs of ovaries (dissected in cold PBS) were homogenized in 500 μl cold PBST at 4.0 meters/second for 20 seconds in a FastPrep-24 Classic homogenizer (MP Biomedicals) using 2 ml Lysing Matrix A tubes (MP Biomedicals). Samples were vortexed, transferred to 1.5-ml tubes and centrifuged at 16,000 g for five minutes at 4°C. One-fifth of the supernatant was collected without disturbing the lipid layer to measure protein content. The remaining supernatant was used to collect the lipid layer and heated for 10 minutes at 70°C in a dry bath. For all experiments (except in Fig 1G), TAG levels were then measured using the Serum Triglyceride Determination Kit (TR0100-1kt, Millipore) according to the manufacturer’s protocol, adapted for 96-well plates. Briefly, 25 μl supernatant from each sample (or 25 μl standards) were transferred in four replicates to 96-well plates, 200 μl TAG reagent was added to half of the wells and Free Glycerol reagent was added to the other half of the wells, and the plate was incubated at 37°C for 30 minutes. Absorbance at 492 nm was measured using a plate reader (Synergy H1, Agilent BioTek). The TAG content was determined by subtracting free glycerol from TAG, and then the fat body TAG content was determined by subtracting ovary pair TAG amounts from whole body TAG amounts. (Note: We did not measure TAG in dissected carcasses directly because fat body cells are often lost during dissection, which introduces technical variability and can lead to inaccurate “per fly” measurements.) For experiment in Fig 1G, a more sensitive Triglyceride Quantification Assay kit (AB65336-1001, Abcam) was used following the manufacturer’s protocol. Briefly, 6 μl sample were used per well in four replicates; 2 μl lipase reagent were added to two replicates and 2 μl lipase buffer were added to the other two replicates. Samples were incubated at room temperature for 20 minutes, after which 50 μl of triglyceride reaction mix were added to all wells and incubated at room temperature for 60 minutes. The absorbance of the plate was read at 570 nm using a plate reader (Synergy H1, Agilent BioTek). Fat body TAG content was determined as described above. Experiments were done in triplicate and statistical analysis was performed using the Student’s *t*-test (GraphPad Prism).

### Glucose, trehalose, and glycogen quantification

For carbohydrate measurements, five seven-day old females (per sample) were washed in PBS and transferred to a 1.5-ml tube. All liquid was removed, and females were homogenized in 100 µl cold PBS (on ice) using a motorized pestle (749521-1500, Kontes). Samples were heated for 10 minutes at 70°C on a heating block and centrifuged at 16,000 g for five minutes at 4°C, and supernatants were collected and stored at −80°C. The Glucose (GO) Assay Kit (GAGO20-1KT, Sigma-Aldrich) was used to measure the amount of free glucose, trehalose (after enzymatic breakdown into two glucose molecules) or glycogen (after enzymatic breakdown into many glucose molecules). For glycogen measurements, 30 µl 1:10 diluted supernatants were transferred to a 96-well plate in six replicates. 100 µl GO reagent containing 2.3 units/ml amyloglucosidase were added to three of the replicates, and 100 µl GO reagent alone were added to the remaining replicates. The plate was incubated at 37°C for 60 minutes, after which reactions were stopped by addition of 100 µl 6 N sulfuric acid. The 540 nm absorbance was measured using a Synergy H1 plate reader (Agilent BioTek), and glycogen levels were calculated based on a glucose standard curve after subtracting the absorbance measured for free glucose in the untreated samples from the absorbance of the samples digested with amyloglucosidase. For trehalose and glucose measurements, 30 µl 1:8 diluted supernatants were transferred to 1.5-ml tubes containing 30 µl Trehalase Buffer (5 mM Tris pH 6.6, 137 mM NaCl, 2.7 mM KCl) with 2.7 units/µl porcine trehalase or 30 µl of Trehalase Buffer alone. Tubes were incubated at 37°C for 24 hours, after which samples were transferred to 96-well plates and processed as described above for glucose measurement using the Glucose (GO) Assay Kit. Free glucose levels (from samples not treated with trehalose) were calculated based on a glucose standard curve. Trehalose levels were calculated based on a glucose standard curve after subtracting the absorbance measured for free glucose in the untreated samples from the absorbance of the samples digested with trehalase. All experiments were done in triplicate and statistical analysis was performed using Student’s *t*-test (GraphPad Prism).

### *Neural Lazarillo* quantitative RT-PCR

Ten adult female fat bodies (scraped off carcasses) or pairs of ovaries were dissected and incubated in RNA*later* Stabilization Solution (ThermoFisher Scientific) for 10 minutes. After RNALater removal, 250 μl lysis buffer from the RNAqueous-4PCR Total RNA Isolation Kit (ThermoFisher Scientific) were added and samples were homogenized using a motorized pestle and RNA extraction proceeded according to the manufacturer’s instructions. Complementary DNA was synthesized from 1 mg total RNA using oligo (dT) primers and SuperScript IV Reverse Transcriptase (ThermoFisher Scientific) according to the manufacturer’s instructions. Complementary DNA was amplified through a 40-cycle reaction (95°C for 3 seconds, 55°C for 3 seconds and 72°C for 20 seconds for *Nlaz*; 95°C for 3 seconds, 55°C for 3 seconds and 72°C for 20 seconds for *RpL32* using previously described primers for *NLaz*, 5′- GGACAACCCTCGAATGTAACT -3′ and 5′- GACGGCGTATGACTCGTAATC -3′ [34], and for *RpL32* (aka *RP-49*; used as a normalization control), 5′- CAGTCGGATCGATATGCTAAGC -3′ and 5′- AATCTCCTTGCGCTTCTTGG -3′ [68]. Mock reactions using water without complementary DNA served as negative controls. Experiments were done in triplicate and statistical analysis was performed using the Student’s *t*-test on the relative *ΔΔ*CT quantification method (GraphPad Prism).

### Western blot analysis

Six fat bodies or pairs of ovaries (per sample) from seven-day-old females maintained on control or high sugar diets were dissected in Grace’s medium and transferred to another well containing Grace’s medium with or without 0.5 μM insulin and incubated for 20 minutes at room temperature. Samples were then transferred to 1.5-ml tubes and 60 μl Protein Lysis Buffer [50 mM Tris-HCl, 0.1% Triton X-100, 200 mM NaCl, 1 mM EDTA, 0.2 mg/mL Sodium Azide, 1:100 Protease Inhibitor Cocktail (P8340, Sigma)] were added. Samples were homogenized with a motorized pestle on ice and centrifuged at 16,000 g for five minutes at 4°C, and supernatants were collected. Protein quantification was performed using a BCA Protein Assay (23225, Pierce) according to the manufacturer’s instructions. 10 µg protein from each sample in Fluorescent Compatible Sample Buffer (Invitrogen) were heated at 95°C in a heating block for 10 minutes prior to loading. Samples were electrophoresed on NuPAGE 4 to 12%, Bis-Tris, 1.5 mm, Mini Protein Gels (Invitrogen WG1401BOX) in NuPAGE MOPS SDS running buffer (Invitrogen) at 200 V for 42 minutes. Samples were then transferred to Low Fluorescence PVDF (Azure) membranes in NuPAGE Transfer Buffer (Life Technologies) at 20 mA for 60 minutes using a Semidry Blotter (ThermoFisher Scientific). Membranes were stained for total protein using Red Protein Stain (Azure) and washed three times in methanol and two times in PBS. Membranes were blocked with AzureSpectra Protein Free Blocking Buffer (Azure) for 60 minutes and incubated with primary antibodies 1:1000 rabbit anti-pAKT [Phospho-Akt (Ser473) antibody #9271, Cell Signaling Technology] and 1:500 mouse anti-*α*-Tubulin (12G10, DHSB) at 4°C overnight. After rinsing twice and washing three times with PBST, the membranes were incubated with 1:10,000 secondary antibodies Azure 700 mouse and 800 rabbit for two hours, rinsed twice and washed three times with PBST, and washed twice with PBS, according to the manufacturer’s protocol. Membranes were imaged using an Azure 600 Western blot imaging system. Quantification of bands and normalization of the bands were made using the BandPeakQuantification macro made by Kenji Ohgane and Hiromasa Yoshioka on FIJI with settings: background width (pixels) 3; estimate background from top/bottom; background estimation by mean.

## ACKNOWLEDGMENTS

We thank the Developmental Studies Hybridoma Bank for antibodies, and the Bloomington Stock Center (National Institutes of Health P400D018537), Vienna Drosophila Resource Center, and Ronald Kühnlein for *Drosophila* stocks. We are grateful to Phil Newmark and his lab members for generously sharing their Azure 600 Western blot imaging system and Synergy H1 plate reader. We also thank members of the Newmark and D.D.-B. labs for helpful discussions during lab meetings. We thank Emily Wessel, Sabi Nagarajan, Ana Caroline Gandara, and Alicia Williams for careful reading of the manuscript and helpful editing suggestions. This work was supported by National Institutes of Health (NIH) grants R01 GM069875 (D.D.-B.), R01 GM125121 (D.D.-B.), and R35 GM140857 (D.D.-B.).

## SUPPLEMENTAL FIGURE LEGENDS

**S1 Fig. Genetically induced obesity has no effect on the maintenance or proliferation of germline stem cells.** (A,B) Line graphs showing average number of germline stem cells (A) or cap cells (B) per germarium at zero, five, 10, and 15 days of adipocyte-specific knockdown of *bmm* or control *LUC*. The numbers of germaria analyzed are shown in parentheses. Data shown as mean±s.e.m. from three independent experiments. ns, no significant difference, two-way ANOVA with interaction. (C,D) Frequencies of EdU-positive (C) or phospho-histone H3 (pHH3)-positive (D) germline stem cells in females at five days of adipocyte-specific knockdown control *LUC* or *brummer*. Numbers of germline stem cells analyzed are shown inside the bars. Data shown as mean±s.e.m. from three independent experiments. ns, no significant difference, Chi-square test.

**S2 Fig. Obesity caused by a high sugar diet is resistant to leanness-inducing genetic manipulation.** (A, B) Adipocytes from seven-day-old females with adult adipocyte-specific *LUC* control RNAi (A) or *brummer* overexpression plus *Lsd2* knockdown (B) on a high sugar diet. Nile red (magenta) labels lipid droplets; Phalloidin (green) outlines cell membranes; DAPI (white) labels nuclei. Images represent projections of three 1-μm confocal sections. Scale bar, 20 μm. (C) Bar graph showing average triglyceride content per female (after subtraction of ovarian triglyceride content) of seven-day-old females with adult adipocyte-specific control *LUC* RNAi, control GFP overexpression, or *brummer* overexpression plus *Lsd2* RNAi knockdown on a high sugar diet. Data shown as mean±s.e.m. from three independent experiments (10 females per genotype per experiment). ns, no significant difference, Student’s *t*-test.

