## Supplementary figures and images for "A high sugar diet, but not obesity, reduces female fertility in *Drosophila melanogaster*"

### Supplemental Figures

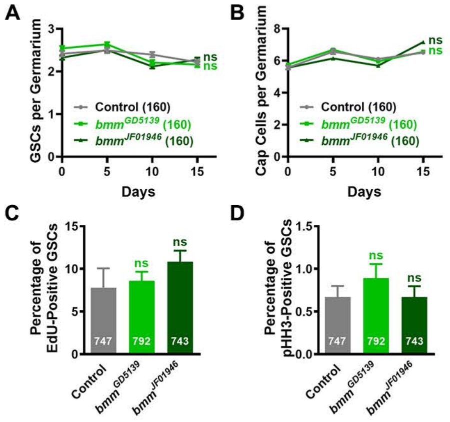


**S1 Fig**
**Nunes & Drummond-Barbosa**

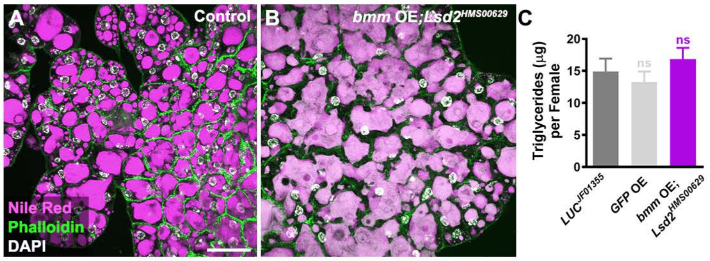


**S2 Fig**
**Nunes & Drummond-Barbosa**
